# BioFeatureFinder: Flexible, unbiased analysis of biological characteristics associated with genomic regions

**DOI:** 10.1101/279612

**Authors:** Felipe E. Ciamponi, Michael T. Lovci, Pedro R. S. Cruz, Katlin B. Massirer

## Abstract

BioFeatureFinder is a novel algorithm which allows analyses of many biological genomic landmarks (including alternatively spliced exons, DNA/RNA-binding protein binding sites, and gene/transcript functional elements, nucleotide content, conservation, k-mers, secondary structure) to identify distinguishing features. BFF uses a flexible underlying model that combines classical statistical tests with Big Data machine-learning strategies. The model is created using thousands of biological characteristics (features) that are used to build a feature map and interpret category labels in genomic ranges. Our results show that BFF is a reliable platform for analyzing large-scale datasets. We evaluated the RNA binding feature map of 110 eCLIP-seq datasets and were able to recover several well-known features from the literature for RNA-binding proteins; we were also able to uncover novel associations. BioFeatureFinder is available at https://github.com/kbmlab/BioFeatureFinder/.

## Background

The emergence of high-throughput sequencing technologies has led to an enormous number of datasets available for researchers, and multiple types of analysis have been built on top of these technologies [1]. It is now possible to use strategies to identify protein binding sites (ex. ChIP [2]/CLIP-seq [3]), alternative splicing (AS) events [4], and differentially expressed genes [5], detect SNPs [6] and achieve a multitude of other applications [7,8], resulting in large sets of genomic coordinates (e.g., binding sites, AS exons, polymorphisms). Such results are particularly challenging to interpret from a biological perspective. Several approaches have been used to characterize genomic coordinates sets and identify the enriched characteristics (features) in these datasets, especially when using the results from ChIP-seq or CLIP-seq experiments [8–12]. However, several of the most commonly used tools in these analyses focus on a particular aspect of the target regions in their process, such as structural models or sequence motifs [9,11,13]. Although these tools provide valuable insight into which characteristics are enriched in the genomic regions associated with these datasets, there is a clear deficiency in the computational tools that can perform more comprehensive analyses and integrate multiple types of sources of variation.

In previous high-throughput studies, some individual features were revealed to be key contributors in large datasets: GC content [14–16], nucleotide composition [14], length [14,16], CpG islands [16,17], conservation [18], microRNA [19,20] and protein-binding [21] target sites, methylation sites [22], single nucleotide polymorphisms (SNPs) [23], microsatellite regions [24] and the aforementioned sequence motifs and structural characteristics of these regions. However, due to biological variability within the genome sequences [25–27], we must also consider that sets of genomic regions are composed of a heterogeneous population of sequences, each with a unique profile of characteristics. While not all these characteristics are enriched and/or important for creating a profile for the whole-region groups, it is possible that a combination of many factors is responsible for separating groups of genomic regions and/or determining the binding of a protein to that particular region.

We present BioFeatureFinder (BFF), a flexible and unbiased algorithm for the discovery of distinguishing biological features associated with groups of genomic regions. In addition, BFF can help discover how these characteristics interact with each other. This is useful for creating an accurate map of the features that are more important for explaining differences between the input genomic regions and the remaining regions of the genome. To achieve this, we applied machine-learning strategies that are already widely used in other transcriptomics, genomics and system biology studies [21,28–31]. Instead of analyzing the genomic regions as individual data points, we analyzed the cumulative distribution functions drawn from the population of regions for each of the features described above and then applied binary classification algorithms to identify which characteristics are more important for group separation. This strategy has already been applied in other studies [32] but is here applied for the first time in the context of classification of features associated with groups of genomic regions on this scale. Furthermore, we aimed to develop a flexible tool that can use data from multiple public databases, such as UCSC GenomeBroswer [33], Ensembl [34], GENCODE [35], ENCODE [36] and others, and is capable of performing in an unsupervised and unbiased way.

## Results and Discussion

For our BFF algorithm, we define “biological features” as the set of characteristics that can be used to distinguish regions from other sections of the genome. These features can include but are not limited to nucleotide content, length, conservation, k-mer occurrence, presence of SNPs, protein-binding sites, microRNA target sites, methylation sites, microsatellite, CpG islands, repeating elements, and protein domains.

Our algorithm can analyze the distribution of values for each feature in a set of genomic regions of interest, compare this distribution to a randomized background to identify which features represent the most distinguishing characteristics associated with the input dataset, and then rank them by importance values. This tool can be used as an important information source for scientists, who can use the data provided to generate new and more accurate hypotheses or guide wet-lab experiments more efficiently. BFF can also be used in large-scale computational projects that can analyze hundreds of datasets with ease and produce consistent results.

For the first time, we apply Big Data strategies in an unbiased way, thereby effectively reducing observer bias, to take advantage of the large amounts of data produced by high-throughput experiments, such as CLIP/CHIP/RNA/DNA-seq, and data deposited in publicly available databases (e.g., UCSC GenomeBroswer [33], Ensembl [34], GENCODE [35], ENCODE [36]) to extract a set of significant information from genomic regions and uncover the latent relationships inside the datasets. First, we present the framework used by BFF in its analytic process, with an overview of the input data types, workflow and output. Second, as a control, we applied BFF to the RBP (RNA-binding protein) RBFOX2 eCLIP-seq (enhanced crosslink immunoprecipitation RNA-sequencing) data because this protein is widely studied, and its binding sites are well characterized in the literature [37–40]. Finally, to showcase potential applications of the algorithm, we analyzed 112 eCLIP datasets obtained from human cell lines that are available in the ENCODE database and identified the biological features associated with the binding sites of all RBPs and their respective importance scores.

### The BioFeatureFinder workflow

BioFeatureFinder focuses on flexibility, consistency and scalability. It is python-based with scalable multi-thread capabilities, is memory-friendly and compatible with most commonly used UNIX-based systems (e.g., CentOS, Ubuntu, openSUSE and macOS) and can be used with a wide range of hardware, from notebooks to HPC clusters. In Figure 1, we show a schematic representation of the BFF workflow. The types of inputs required to use the algorithm are as follows: a set of BED coordinates with genomic regions of interest (e.g., CLIP/CHIP-seq binding sites, promoter regions for differentially expressed genes, splice sites for alternatively spliced exons/introns and others); and compatible fasta files with sequences (e.g., Reference genome transcriptome). However, for increased accuracy, it is also possible to use a GTF/GFF file with region annotations (exons, introns, CDS, UTR and others). These can be optionally used to increase the number of features that are analyzed using BED files with genomic regions of biological features (e.g., microRNA sites, methylation sites, CpG islands, protein binding sites, SNPs and mutations, repeating elements and multiple bigWig files with phastCons scores for multiple alignment).

**Figure 1:**
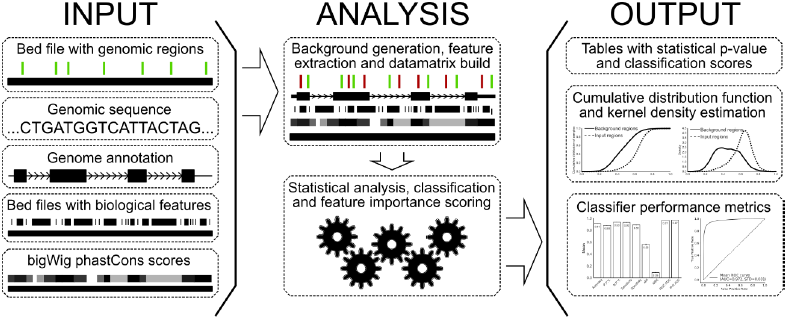
Schematic overview of BioFeatureFinder workflow. Schematic representation of the 3 steps involved in the BioFeatureFinder workflow. The first step (INPUT) involves the selection of genomic regions of interest to analyze and the biological information to be extracted from these regions (such as nucleotide sequence or bed files with biological data). The second step (ANALYSIS) refers to the process by which BFF converts the biological information for genomic regions in a numerical datamatrix and then proceeds to analyze this information using both statistical and machine-learning approaches. The final step (OUTPUT) represents the information that is extracted from our algorithm, comprising of both easily human-readable tables and graphical representations of both the biological features and the overall classifier performance.

The analytical process of the algorithm is divided into two sub-sections: Build datamatrix and Analyze features. Building the datamatrix starts with selecting an appropriate background to compare the input regions of interest, which are obtained using the *shuffle* function of bedtools. Although it is not required, the use of a reference annotation improves the algorithm’s accuracy by guiding the included/excluded background regions. The total number of background regions (B) is proportional to the number of regions in the input list (I) of bed coordinates, which can be represented by the following formula: *B* = *I* * *N*, where N is an integer variable that can be set as an option (default = 3; i.e., the number of background regions is 3 times the number of input regions). These two sets of regions are then used to produce a datamatrix. Each bed entry in the regions is converted into a line in the matrix, and each feature corresponds to a column. Every feature is represented as numeric value, which can be a continuous, discrete or Boolean variable. To obtain these values, BFF uses multiple freely available tools such as bedtools [41] for the nucleotide content and counting intersections with features in bed format, bigWigAverageOverBed [42] for extracting conservation scores, EMBOSS wordcount [43] for k-mer counting, Vienna’s RNAfold [44] for RNA secondary structure MFE (minimum free energy) values and QRGS Mapper [45] for G-quadruplex scoring. Designed with a modular concept, new functions and sources of data can be easily added by researchers to answer project-specific questions.

Once the matrix is created, the algorithm runs a two-step analysis to identify important features in the dataset. The first step is to analyze each feature in the matrix with a two-sample Kolmogorov-Smirnov test (KST), with implementation by SciPy [46], to compare the distribution of values of the regions of interest (group 1) to background regions (group 0). In addition to using a statistical tool to identify the significant features, it is possible to use KST as a tool for feature selection by extracting statistically significant features that are correlated with differences between the groups, a strategy that has been shown to improve the classification performance of high-dimensional data [47–49]. As an additional benefit, filtering the features by KST p-values reduces the size of the datamatrix used in the following classification step, which can be helpful for reducing both the computational time and the resources used in the analyses. The second step involves the use of a Stochastic GradientBoost Classifier (St-GBCLF) from Scikit-learn [50] that can naturally handle mixed datatypes, is fairly robust to outliers and possesses reliable predicative power. Additionally, this method has been shown to be preferable for high-dimensional two-class prediction [51,52]. Additionally, similar to other ensemble methods, St-GBCLF is less likely to suffer from overfitting [53]. This stage will use the feature values extracted from the matrix (which can be filtered, or not, by KST) for each group (0, background and 1, input) and calculate the feature importance, a score that measures how valuable each feature is in the decision-making process of the trees [54]. Higher importance values indicate that the feature is considered in key decisions and can thus be inferred to have more biological significance. To address the class imbalance problems inherent to these types of analysis [52,55], our algorithm draws a random sample from the background (group 0), which is the same size as the input dataset (group 1), thereby increasing the overall accuracy of the classifier. However, to address biological variability, our algorithm performs multiple classification runs, drawing a new sample of background regions each time. The final classification score is calculated as the average of the importance values obtained in each classification run.

Both classification and statistical results are compiled into a table, which allows easy interpretation. Additionally, graphical representations of each feature are output in both the cumulative distribution function (CDF) and kernel density estimation (KDE) plots. This allows the visualization of the distributions found in the input and background and leads to conclusions about how the distribution is shifted from the reference. Additionally, classifier importance and KS test values are output in bar charts. Finally, classifier performance is measured by several parameters: accuracy, sensitivity, sensibility, positive predictive value, negative predictive value, adjusted mutual information (aMI), mean squared error (MSE) value, receiver operating characteristic (ROC) and precision-recall (P-R) area under curve. These metrics are output in both table (with scores) and graphical (bar charts and curves) format. Together, these outputs can be used to explore the data and identify features that may be of significance in a biological context.

### Analysis of RBFOX2 eCLIP dataset

To evaluate the performance of our algorithm, we analyzed the subset of RNAs binding regions bound by the RNA binding protein RBFOX2. We used public eCLIP experiments deposited in the ENCODE database. Within this set, we found 922 statistically significant biological features that were used in the classification step (Figure 2A). The classifier provided an overall average of accurate predictions of 9 out of 10 times. The scores were 91% mean accuracy score, 88% positive predictive value (P.P.V.), 93% negative predictive value (N.P.V.), 93% sensitivity, and 89% specificity. We also obtained average scores for adjusted mutual information (aMI) and mean squared error (MSE) of 0.56 and 0.09, respectively. Both the receiver operating characteristic (ROC) and Precision-Recall (P-R) area under curve (AUC) were measured at 0.97 (Figure 2A, Additional File 1). Among all of the statically significant features, our approach identified 16 features with a relative importance score of at least 10% (i.e., 1/10 of the importance score of the highest scoring feature, Figure 2B, Additional File 2), rediscovering known literature-reported features and novel features. As a known feature, we found conservation of the binding site and enrichment of the canonical GCAUG k-mer; and within novel features, we found higher GC-content in the binding sites, lower MFE for RNA secondary structure, a higher G-quadruplex score and overlap of binding sites with DDX6 and other RBPs.

**Figure 2:**
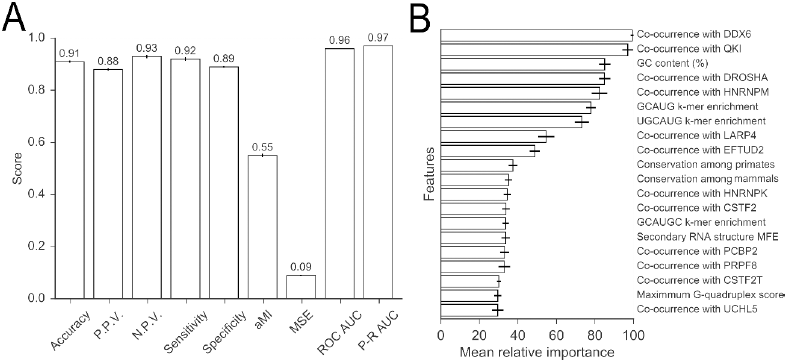
BioFeatureFinder accurately identifies biological features associated with RBFOX2 CLIP-seq binding sites. **A.** Classifier performance scores as the mean values for RBFOX2 eCLIP sites, represented as a bar chart (P.P.V.: Positive predictive value, N.P.V.: Negative predictive value, aMI: Adjusted mutual information, MSE: Mean squared error, ROC AUC: Receiver operating characteristic area under curve, P-R AUC: Precision-Recall area under curve).; **B.** Feature importance score (white) and Kolmogorov-Smirnov test value (KS-test in gray) for the top 20 features associated with RBFOX2 sites, represented as horizontal bar chart. Black vertical bars represent the standard deviation found for each scoring parameter.

We identified enrichment of the GCAUG 5-mer as a major feature that characterized the RBFOX2 eCLIP binding sites by both KS test (p-value < 0.001) and variable importance in classification. Our analysis indicated that 33% of the binding sites identified in the RBFOX2 eCLIP contained at least 1 repetition of the GCAUG motif, whereas only 6% randomized background regions exhibited at least 1 instance of this motif (Figure 3A). Additionally, we found significant enrichment of the UGCAUG 6-mer, which occurred in 22% of binding sites, in contrast to 2% of the background regions (Figure 3B). Both results are consistent with the RBFOX2 nucleotide sequence motif enrichment and occurrence in RNA binding sites [37,40], indicating that our algorithm successfully recovered known features.

**Figure 3:**
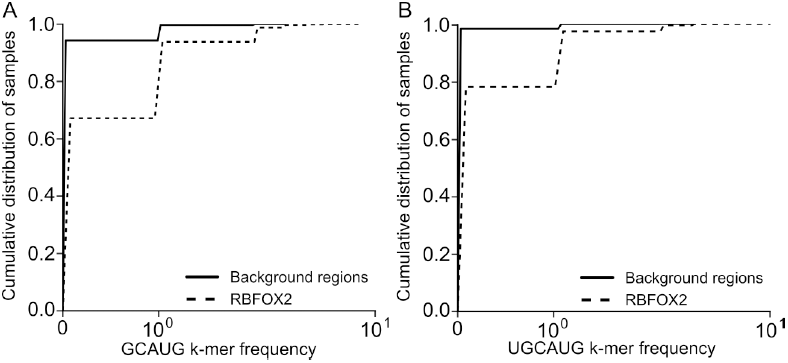
RBFOX2 CLIP-seq binding sites are enriched for the (U)GCAUG motif. A-B. Cumulative distribution function curves for GCAUG (A) and UGCAUG (B) k-mer sequences. Y-axis shows the cumulative distribution of samples, and X-axis indicates the number of occurrences for each k-mer. Solid lines represent randomized background regions, and dashed lines represent RBFOX2 binding sites

Interestingly, we identified several major components associated with the RNA secondary structure of the RBFOX2 binding sites. GC content had the second highest importance value of all features, with RBFOX2 binding sites exhibiting a higher distribution of GC than randomized background regions and most of the binding sites having a range of 50% to 80% GC content in their sequences, whereas the background regions were more evenly distributed (between 20% to 60% -Figure 4A). It is known that RNA regions with higher GC content correlate to more stable secondary RNA structures [56], with alterations in splicing patterns through an effect on pre-mRNA secondary structure [57], a known mechanism for RBFOX2 splicing regulation [37]. Although the importance of RNA secondary structure as a guiding factor for RBP binding was shown before [13,58,59], our data pointed to RBFOX2 because we identified that the minimum free energy (MFE) for RNA folding is a major feature for distinguishing protein binding sites from a randomized background. We identified 70% of binding sites with an MFE lower than 0, indicating the possible existence of a localized secondary structure, while only 47% of background regions exhibited similar behavior (Figure 4B). Additionally, we also identified the presence of G-quadruplexes, a specific type of secondary structure, as enriched in RBFOX2 binding sites. Our analysis indicates that 53% of RBFOX2 binding sites had a positive score for their presence, in contrast to only 15% of background regions (Figure 4C).

**Figure 4:**
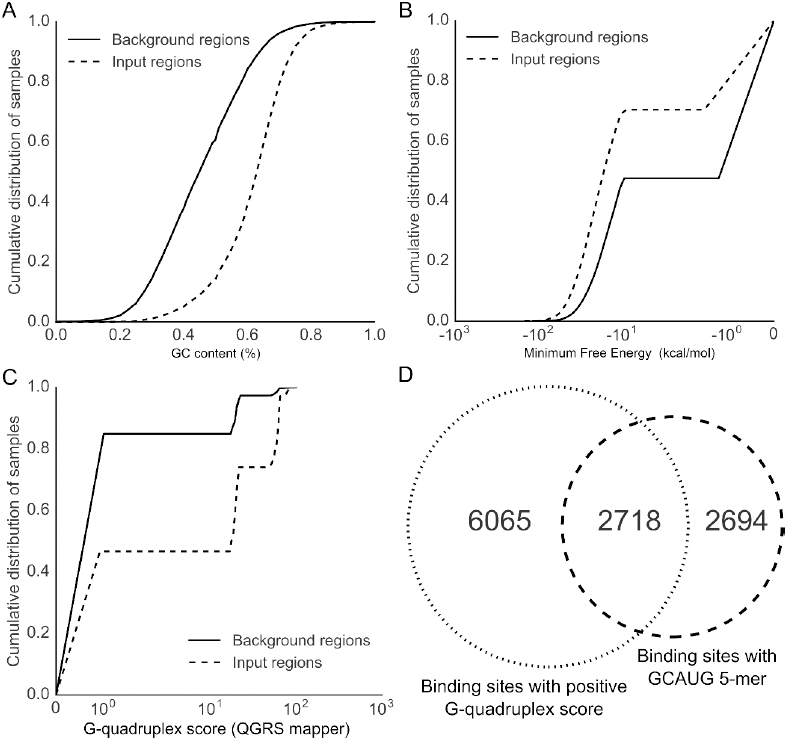
RBFOX2 binding regions are enriched in GC content within structured RNA regions A-C. Cumulative distribution function curves for GC content (A), Minimum Free Energy (MFE, B) and maximum G-quadruplex score (C). Y-axis shows cumulative distribution of samples and X-axis indicates values obtained for each feature. Solid lines represent randomized background regions, and dashed lines represent RBFOX2 binding sites. **D**. Two-way Venn diagram showing the overlap between number of peaks identified with the GCAUG k-mer and those with a positive G-quadruplex score.

This result is particularly interesting because although this feature was not previously associated with RBFOX2, it can be supported extensively by literature evidence. First, RBFOX2 has been described as a member of the RG/RGG family of RNA-binding proteins, with an RNA recognition region rich in arginine-glycine rich regions that is known as RGG-box [60]. Second, other proteins from this family were shown to bind to RNA G-quadruplex by their RG/RGG regions [61]. Third, the existence of G-quadruplexes in intronic regions can have impacts on alternative splicing regulation [62,63]. Taken together, these results indicate that secondary RNA structure may play a larger role in RBFOX2 targeting for binding sites than previously assumed, combined with the existence of the GCAUG motif for increased accuracy in target selection. This is further evidenced by the fact that 50% of the binding sites containing GCAUG were also positive for the presence of G-quadruplexes (Figure 4D).

Finally, the most important feature, an association previously unreported in the literature, represents the overlap of RBFOX2 binding sites with DDX6 binding sites, an RNA helicase. We identified 18% of RBFOX2 peaks with at least 1 nucleotide position in common with DDX6 binding sites, which is significantly higher than the value obtained for randomized background regions that scored almost 0% of overlap (Figure 5A). Although this association is novel, it is known that DDX6 homologs in *S. cerevisiae* interacts with EFTUD2 homolog, another RBP identified as an important feature for RBFOX2, which suggests that the interaction between these proteins is evolutionarily conserved. [64]. We also identified several other RPBs with significant overlap with RBFOX2, including known splicing regulators and/or components of the spliceosome such as HNRNPM, EFTUD2, PRPF8, QKI, HNRNPK and PCBP2 (Additional File 2).

**Figure 5:**
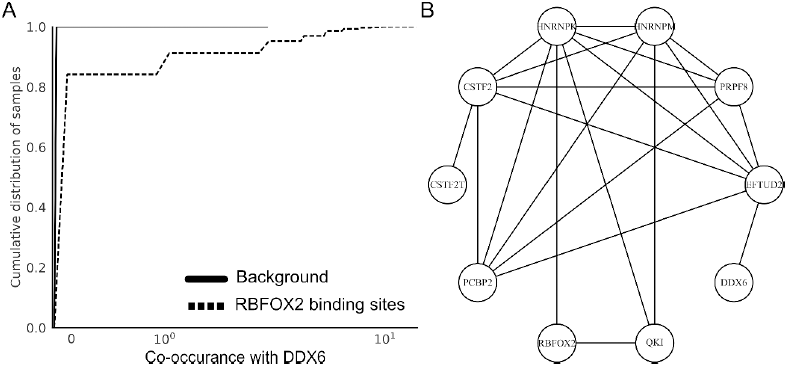
Discovered features include proteins with overlapping RNA binding profile and connected protein-protein interaction (PPI) network. **A.** RNA binding protein DDX6 was featured as having overlapped binding sites with RBFOX2. Cumulative distribution function curves where Y-axis shows the cumulative distribution of samples and X-axis indicates number of occurrences per overlap. Solid lines represent randomized background regions, while dashed lines represent RBFOX2 binding sites. **B.** PPI network created based on protein interactions obtained from BioGrid 3.4 and StringDB 10.5. Each node represents a different RBP, and lines represent known and inferred protein-protein interactions between them.

Using data available from BioGrid 3.4 [65] and STRINGdb 10.5 [66], we found that these targets were associated with RBFOX2 through a curated protein-protein interaction network (Figure 5B), with RBFOX2 directly interacting with QKI [67–70] and HNRNPK [70]. The latter, in turn, has been shown to interact with EFTUD2 [71].

Taken together, our results indicate the existence of a combinatorial mechanism of both RNA structure and nucleotide sequence to direct binding specificity to RBFOX2. However, our analysis is limited to identifying the similar and diverging characteristics of their binding sites. For a deeper understanding of the relationship between RBFOX2 binding features, further experiments are required. They would contribute to the comprehension of target binding specificity and would shed light on novel biological functions for RBFOX2.

### Identification of important features for 110 RNA-binding protein-binding sites from ENCODE

To showcase the potential applications of BioFeatureFinder in high-throughput studies, we applied our algorithm to the large dataset of 110 eCLIP-seq available at ENCODE. First, we identified the preferential binding regions for each RNA Binding Protein (RBP) in the dataset (Figure 6, Additional File 6), with our results indicating that 59% of the analyzed proteins had preferential binding to the intronic regions. The second most frequent region was 3’UTR (15%), followed by CDS (15%), 3’ splice site (7%), 5’ splice site (2%) and the 5’UTR (2%). Among the identified preferential regions for the RBPs, some were already known from the literature (such as U2AF1 [72], U2AF2 [72], SF3A3 [73], PRPF8 [74], FMR1 [75], PUM2 [76], TIA1 [77], TARDBP [78] and RBFOX2 [37]), which demonstrates that our algorithm correctly identified their binding region preferences. We used this information to generate the appropriate background for each RBP. Overall, our algorithm performed consistently, with an average accuracy of 0.9 and average ROC and Precision-Recall AUCs of 0.95. The largest variance was obtained for the aMI (adjusted mutual information) scores, with an average of 0.57 and standard deviation of 0.16 (Figure 6B, Additional File 2). We also observed a strong correlation (Pearson’s R^²^ ≥ 0.95) between aMI (Adjusted Mutual Information, Figure 7A) and MSE (Mean Squared Error, Figure 7B) scores with overall classifier Accuracy, indicating that RBPs with higher aMI scores tended to reach a higher degree of resolution of the binding site features. This can be inferred to be a consequence of the binding characteristics of the RBPs. For example, TARDBP has a strict set of characteristics that guide their binding to specific targets, whereas other RBPs, such as SF3B1, appear to have a higher degree of flexibility in their binding target selection (Additional File 2).

**Figure 6:**
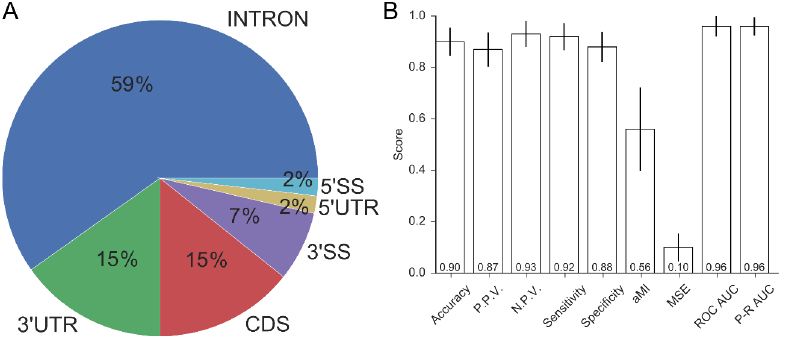
BioFeatureFinder performs consistently and accurately for 110 RBPs that bind to multiple transcript regions. **A.** Pie chart showing the percentage of RBPs with preferential binding to each transcriptomic region. Each slice of the chart corresponds to a different region (Intron, 3’UTR, CDS, 3’SpliceSite, 5’UTR, 5’SpliceSite) and percentages correspond to number of RBPs with more binding sites to that region. **B.** Classifier performance scores as the mean values for 110 eCLIP sites, shown as bar charts (P.P.V.: Positive predictive value, N.P.V.: Negative predictive value, aMI: Adjusted mutual information, MSE: Mean squared error, ROC AUC: Receiver operating characteristic area under curve, P-R AUC: Precision-Recall area under curve). Black bars represent the standard deviation found for each scoring parameter

**Figure 7:**
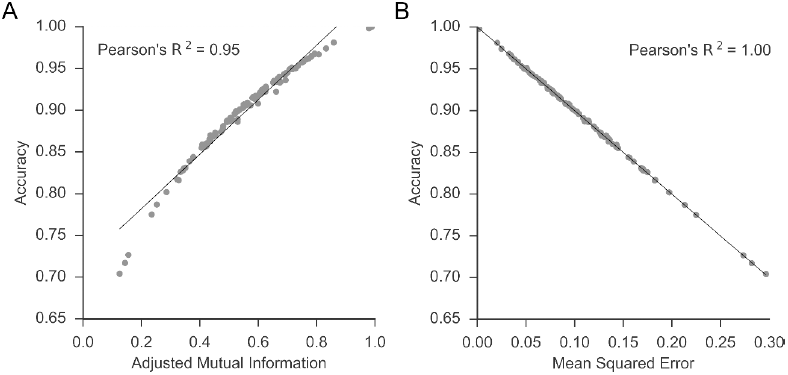
Classifier overall accuracy significantly improves for RBPs with strict set of characteristics defining their binding sites. **A-B.** Scatter plot showing relationship observed between Accuracy scores (Y-axis) and aMI (A) and MSE (B, X-axis). In both cases, a high degree of correlation was identified by Pearson’s R^²^ (>= 0.95).

Overall, we identified three major classes of features that were important for the determination of binding site selection for this group of RBPs: K-mer enrichment (motifs), secondary RNA structure and overlap with other RBPs (Figure 8A, Additional File 7). All 110 RBPs have at least one of these as an important feature for the classification of their binding site with 10% or more relative importance. Furthermore, we found that most of the RBPs (56%) have a combination of these three factors as important features for the determination of binding site specificity. We identified 107 (out of 110) RBPs with some degree of overlap with at least 1 other RBP, which reflects the characteristic of RBPs working in protein complexes to perform biological functions [79,80]. We recovered information for known protein complexes, such as FMR1-FXR1-FXR2 [68,81] (Figure 8B), identifying 68% of FXR1 binding sites with overlap with FXR2 binding sites and 58% with overlap with FMR1 binding sites. Interestingly, the reciprocal did not hold true, with only 19% of FMR1 and 22% of FXR2 binding sites having an overlap with FXR1 (see Additional File 4), which might reflect the molecular dynamics involved in the formation of the complex [81]. In addition, we identified overlaps in the binding sites of RBPs without any previous association reported, such as the AGGF1, which had 40% overlap with GTF2F1, (see Additional File 4). While this information may indicate that they only bind to the same targets in similar positions, it could also suggest the existence of some biological relationship between proteins that is yet to be uncovered.

**Figure 8:**
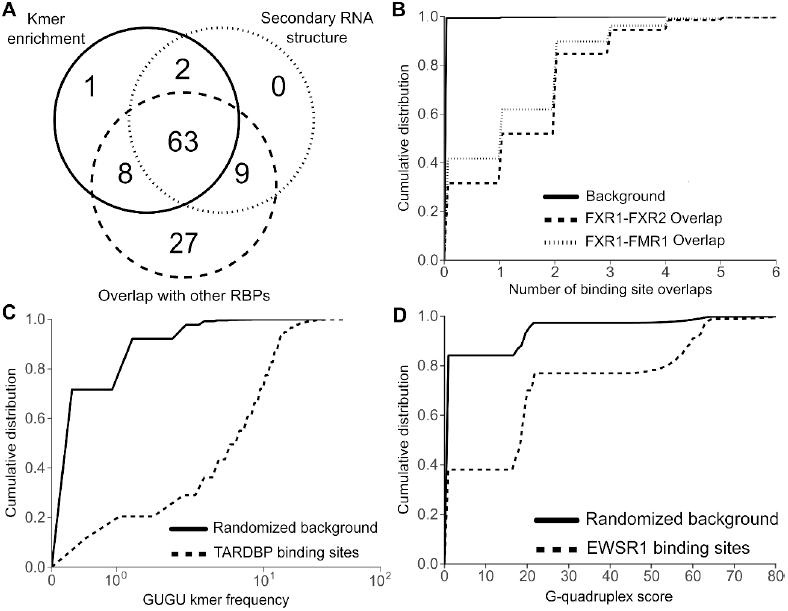
RNA-target selection by RNA-binding proteins is a multi-factorial biological process requiring cis-and trans-regulatory factors. **(A)** Three-way Venn diagram showing overlap between RBPs identified with at least 1 K-mer enrichment (solid line), secondary RNA structure (dotted) and overlap of binding site with other RBPs (dashed) as an important feature for the characterization and group classification of their binding sites. **B-D.** Cumulative distribution function curves for FXR1 binding site overlaps with FMR1 and FXR2 (B), GUGU k-mer enrichment in ARDBP binding sites (C) and EWSR1-binding site maximum G-quadruplex score (D). Y-axis shows the cumulative distribution of samples, and X-axis indicates the values obtained for each feature. The solid lines represent randomized background regions, and dashed lines represent RBPs binding sites.

Analysis of K-mer enrichment revealed that 74 RBPs had at least 1 K-mer with 10% or more relative importance (see Additional File 7); although this represents the majority of analyzed RBPs (66%), it also shows that a nucleotide sequence is not a requirement for directing the binding of all RBPs to their targets. We managed to recover several well-known examples from the literature, such as TARDBP’s GUGU repeats [82], which are present in 88% of binding sites (Figure 8C). Other known examples include QKI [83] (ACUAA in 57% and UAAC in 68% of binding sites), PUM2 [75] (UGUA in 73% of binding sites), PTBP1 [84] (UCUU, 80%), HNRNPC [85] (UUUU, 63%), HNRNPK [86] (CCCC, 87%), KHDRBS1 [87] (UAAA, 80%) and TIA1 [88] (UUUU, 42%). In addition, our algorithm identified motifs for other 47 RBPs, which were found in approximately 30% of binding sites and had an at least 15% difference compared to the background (Table 1, see Additional File 3).

**Table 1.** Nucleotide motifs identified by BioFeatureFinder for 48 RNA-binding proteins. RBPs with nucleotide sequences (motifs) identified as important features were analyzed for percentage of binding sites (BS), which had the identified motif compared to the amount sampled from the background (BG). Differences (Diff) in percentage points (Diff = BS% – BG%) are also represented.

| RBP | Motif | Binding sites (%) | Background (%) | Difference |
| --- | --- | --- | --- | --- |
| FKBP4 | GGGG | 72 | 17 | 55 |
| DDX42 | GGGG | 70 | 17 | 53 |
| NKRF | GGGG | 71 | 19 | 52 |
| XRN2 | GGGG | 70 | 18 | 52 |
| TRA2A | GAAGA | 60 | 11 | 49 |
| DDX59 | CCCC | 67 | 19 | 48 |
| SRSF7 | GUGUG | 53 | 7 | 46 |
| PCBP2 | CCCU | 73 | 26 | 47 |
| GRSF1 | GGGG | 65 | 18 | 47 |
| EIF4G2 | GUGUG | 52 | 7 | 45 |
| SRSF9 | GGAG | 74 | 32 | 42 |
| SERBP1 | CGCC | 54 | 12 | 42 |
| KHSRP | UUGU | 69 | 26 | 43 |
| SLTM | GGGC | 62 | 20 | 42 |
| FUBP3 | UUGU | 66 | 25 | 41 |
| AKAP8L | GGGG | 59 | 18 | 41 |
| GEMIN5 | GCCG | 48 | 9 | 39 |
| FASTKD2 | GGGG | 54 | 17 | 37 |
| DKC1 | GUGUG | 42 | 4 | 38 |
| SRSF1 | GGAG | 69 | 32 | 37 |
| TAF15 | GAGG | 64 | 29 | 35 |
| DDX3X | GCGG | 54 | 22 | 32 |
| HNRNPM | UGUG | 57 | 25 | 32 |
| HNRNPA1 | UUAG | 49 | 17 | 32 |
| SUPV3L1 | GGGG | 46 | 15 | 31 |
| ZRANB2 | GGUG | 50 | 20 | 30 |
| CPSF6 | GAAGA | 41 | 12 | 29 |
| AARS | GGGG | 44 | 16 | 28 |
| RPS11 | GCGG | 32 | 3 | 29 |
| U2AF2 | UUUC | 61 | 33 | 28 |
| HNRNPU | GGGG | 44 | 16 | 28 |
| AGGF1 | CACAC | 33 | 6 | 27 |
| DDX6 | GGGG | 42 | 17 | 25 |
| HLTF | GAAA | 50 | 26 | 24 |
| SAFB2 | GAAG | 50 | 27 | 23 |
| SFPQ | UGUG | 51 | 28 | 23 |
| CDC40 | GGGG | 40 | 18 | 22 |
| SUGP2 | UCUU | 49 | 27 | 22 |
| SUB1 | UGUG | 44 | 22 | 22 |
| DGCR8 | GGGG | 32 | 11 | 21 |
| XRCC6 | CUGG | 50 | 29 | 21 |
| PRPF8 | AGGU | 47 | 26 | 21 |
| SLBP | GAGC | 32 | 12 | 20 |
| LSM11 | GCUG | 43 | 24 | 19 |
| NONO | UGUG | 40 | 24 | 16 |
| TROVE2 | UUGACU | 16 | 1 | 15 |
| SF3B1 | GGGG | 24 | 9 | 15 |

Finally, our algorithm also identified 74 RBPs with secondary RNA structure (either by lower Minimum Free Energy, MFE, calculated by Vienna’s RNAfold, or by a higher G-quadruplex score, from QGRS Mapper) as an important feature for classification (see Additional File 7). As an example, the increased G-quadruplex score for EWSR1 binding sites, which is a known RBP that binds to these types of RNA structures [89], with 62% of the sites exhibiting a positive score, whereas only 16% of the background regions showed the same behavior (Figure 8D). Another RBP that we identified as a binding secondary RNA structure is XRN2, which had 86% of binding sites with an MFE score lower than 0, whereas only 51% of background regions had values lower than 0. This particular RBP is reported to bind R-loop structures formed by G-rich pause sites associated with transcription termination [90], in accordance with our findings for motif enrichment, as we found that XRN2 had 70% of its binding sites containing a GGGG 4-mer, while only 18% of its background regions had the same 4-mer (Table 1). Other known examples from the literature that we recovered include: FMR1, also known to bind to G-quadruplexes [61,91] and DDX3X, DDX6, DDX24 and DHX30, which are RNA-helicases. In addition, we also identified 34 RBPs with positive G-quadruplexes and 15 percentage points or more of difference compared to the background. Of these, 20 RBPs also exhibited an enrichment of GG repeats in their binding sites (Figure 9, Table 1, Additional File 5), which is a known characteristic for these structures [92]. They are: AARS, AKAP8L, CDC40, DDX3X, DDX42, DDX6, DGCR8, FASTKD2, FKBP4, GRSF1, HNRNP, NKRF, SLTM, SRSF1, SRSF9, SUPV3L1, TAF15, XRCC6, XRN2, and ZRANB2.

**Figure 9:**
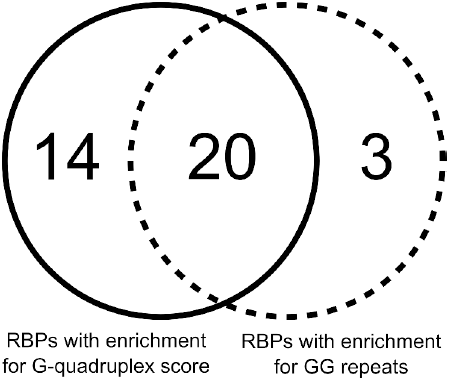
RBPs with enrichment of GG repeats in their motifs also have higher G-quadruplex scores. Two-way Venn diagram showing the overlap between number of RBPs identified with GG repeats in their enriched K-mers (dashed) and RBPs with positive maximum G-quadruplex score (solid) as important features for group classification.

## Conclusion

Our results show that BioFeatureFinder is an accurate, flexible and reliable analysis platform for large-scale datasets while simultaneously providing a method to control observer bias and uncover latent relationships in biological datasets. By considering each genomic landmark as a separate data point in a distribution, we developed a novel implementation. This method combines statistical analysis and Big Data machine-learning approaches to provide accurate representations of the differences in sets of genomic regions and identify the characteristics that contribute more to the separation of these groups.

As demonstrated by our analysis of the RBFOX2 dataset, our algorithm managed to recover multiple characteristics known from the literature, including nucleotide sequences for binding motifs, and infer protein-protein interaction from the overlaps between binding sites. In addition, we uncovered new associations that might link RBFOX2 to the targeting of specific RNA secondary structures to increase RBFOX2 binding specificity, a hypothesis that is strengthened by inferences from the literature from multiple sources.

Furthermore, our analysis of 110 RNA-binding proteins’ CLIP-seq data from ENCODE recovered several well-known features from the literature, including major characteristics that influence the targeting of these proteins. The results for this dataset indicate RNA-target selection by RNA-binding proteins as a multi-factorial mechanism, which demands the existence of both cis-and trans-regulatory factors to increase the RBP affinity to the target site. Among those features are important factors, such as the existence of a particular set of nucleotide sequences (binding motif); accessibility of the target site via RNA secondary structure; and the neighboring RBP context (i.e., other proteins binding to neighboring/same region), which all contribute to determining whether a particular RBP will bind to its target site. Additionally, our results suggest that the binding of RBPs to targets is heavily dependent on the cellular context, with some proteins relying on fewer features for directing their binding specificity (i.e., the presence of a sequence is sufficient for recognition by the RBP), while other proteins require a more complex targeting context, with multiple features involved in the binding of the RBP (i.e., requiring a specific sequence, nearby accessory proteins and a specific RNA structure). Together, our results not only deepen our knowledge of how these proteins select their targets in a broader scenario but also demonstrate how our approach can be applied to large-scale datasets from high-throughput experiments with a high degree of reproducibility.

Although the present study focused on the applications of BioFeatureFinder for RNA-binding proteins, our algorithm can be applied to any type of genomic landmark. Some examples of regions that could be analyzed using BFF include: splicing sites for alternatively spliced exons (or whole exons), target sites for microRNAs, binding sites for DNA-binding proteins (for example, ChIP-seq data), promoter regions from differentially expressed genes, microsatellite/genomic markers, SNPs, whole transcript regions (5’UTR, 3’UTR and CDS) and any type of dataset that can be converted into a BED format.

## Materials and methods

### Extraction of information about biological features and RNA-binding proteins binding sites

We downloaded tracks from biological features associated with genomic features from the UCSC Genome Browser [33] for the human genome hg19, downloading tracks for conservation scores (phastCon scores in bigWig format), benign and pathological CNVs, common and flagged SNPs, TS microRNA target sites, CpG islands, layered H3K4Me1/3 and H3K27Ac and microsatellites. Additionally, we obtained data for 112 RNA-binding proteins (RBP) available from ENCODE [36] eCLIP experiments (Additional File 8: Table S1), downloading the bed files containing the narrowPeaks obtained for hg19 (Additional File 8: Table S2). Additionally, our algorithm is integrated with BedTools [41] (intersect, getfasta and nuc functions), UCSC’s bigWigAverageOverBed [42], EMBOSS wordcount [43], Vienna’s RNAfold [44] and QRGS Mapper [45]. Unless otherwise stated, all tracks were either downloaded or converted into BED format. We used GENCODE’s [35] GRCh37.p13 as a reference genomic sequence, along with release 19 of the comprehensive annotation.

### Preferential region identification and background selection

To identify preferential binding regions for each dataset analyzed, we separated the transcripts into 6 major regions: 5’UTR, 3’UTR, CDS, introns, 5’ splice sites and 3’ splice sites. We then used bedtools intersect to count how many RBP binding sites occurred in each region. These values were normalized by Z-score, and the highest scoring was selected as the preferential binding region. A randomized background was generated using bedtools shuffle (with -excl, -incl, -chrom and -noOverlapping options), using the GTF reference containing the preferential binding region (or regions) to guide the selection of regions, excluding overlaps with input regions (binding sites) and other randomized background regions. For RBPs with small differences in binding site z-scores (less than 10%) in their highest-scoring regions, we selected the 2 highest scoring regions and used both as references for background generation. For each RBP dataset, we generate a randomized background region with three times the number of input regions (binding sites).

### Assembly of a data matrix with biological features

The genomic regions (binding sites and background) and their associated biological features are converted to a numerical matrix, where each line is one region and each column is a biological features associated with that position. To convert biological features in numerical data, we used several freely available software. For most features, we use BedTools intersect (-s and -c options) to count the number of occurrences of that feature in the corresponding region. To obtain nucleotide sequence information, we used a combination of BedTools getfasta and nucBed (both with -s option). For the conservation score we used the tool bigWigAverageOverBed to obtain the average conservation score of covered bases in the region. For k-mer analysis, we used EMBOSS wordcount to count the number of occurrences of each 4-mer, 5-mer and 6-mer in each region. For RNA structure analysis, we used both Vienna’s RNAfold to calculate the lowest possible MFE (with -g option) and QGRS Mapper to calculate the maximum non-overlapping G-quadruplex score for each region. All operations were performed while considering strandedness.

### Group selection and statistical analysis of features

For analysis, the input and background regions are separated into groups (1 and 0, respectively) using the unique identifier created during the datamatrix assembly and stored as the “name” field in the generated BED file. For all features in the matrix, we performed a Kolmogorov-Smirnov test by comparing the cumulative distribution function of the input regions (group 1) to the background (group 0) to filter out non-significant features between the groups. Features with a q-value ≤ 0.05, as adjusted by Bonferroni, were selected for further analysis using the classification algorithm. This filtering aims to provide significantly different features between the groups and to minimize the noise introduced by non-significant features while simultaneously reducing the computational time required for the classification step and the overall required time for analysis.

### Classifier and feature importance estimation

To evaluate the importance of each feature’s ability in separating the genomic region groups, we chose to use a stochastic gradient boost classifier python implementation from Scikit-learn [50]. The classifier was used with the following parameters: number of estimators = ‘1000’; learning rate = ‘0.01’; max depth = ‘8’; loss = ‘deviance’; max features = ‘sqrt’; minimum number of samples per leaf = ‘0.001’; minimum number of samples to split = ‘0.01’; random state = ‘1’ and subsample = ‘0.8’. Importance values for each feature are calculated at every run, with the final value representing the mean scores and their corresponding standard deviation. The same scoring strategy is employed for relative importance score (percentage relative to the most important feature), accuracy, positive predictive value, negative predictive value, sensitivity, sensibility, ROC and Precision-Recall AUCs.

### Availability

The BioFeatureFinder software is available for download at GitHub [93] and is included as Additional file 9 for archival purposes.

### Abbreviations

aMI: Adjusted mutual information
AUC: Area under curve
BFF: BioFeatureFinder
CDF: Cumulative distribution function
CDS: Coding sequence
ChIP-seq: Chromatin Immunoprecipitation sequencing
CLIP-seq: Cross-linking Immunoprecipitation sequencing
CpG: 5’—C—phosphate—G—3’
DNA: Deoxyribonucleic acid
eCLIP-seq: enhanced Cross-link Immunoprecipitation sequencing
FTD: Frontotemporal dementia
GFF: General feature format
GTF: General transfer format
KDE: Kernel density estimation
KST: Kolmogorv-Smirnov Test
MFE: Minimum free energy
MSE: Mean squared error
N.P.V.: Negative predictive value
P.P.V.: Positive predictive value
P-R AUC: Precision-Recall area under curve
RBP: RNA-binding protein
RNA: Ribonucleic acid
ROC AUC: Receiver operating characteristic area under curve
SNP: Single-nucleotide polymorphism
St-GBCLF: Stochastic gradient boost classifier
UTR: Untranslated region

## Declarations

### Acknowledgments

Funding was received from the São Paulo Research Foundation (FAPESP grants 12/00195-3, 13/50724-5). Felipe E. Ciamponi was recipient of a FAPESP fellowship (14/25758-6, 15/25134-5, 16/25521-1). Dr Mario H. Bengtson and Dr Marcelo Brandão who provided valuable scientific insights.

### Competing interests

The authors declare they have no competing interest.

### Data availability

All data used in this study is publicly available from ENCODE database and the developed software and raw results are available on GitHub.

### Author contribution

FEC, MTL and KBM conceived the project and designed its overall goals. FEC and MTL designed the algorithm and wrote the Python code. FEC implemented classification algorithms for biological datasets and performed the analysis of the eCLIP datasets. PRSC assisted in developing, debugging, testing and benchmarking the algorithm. MTL and KBM coordinated the project. FEC wrote the manuscript. All authors read and approved the final manuscript.

### Supplementary material

Additional File 1: Deviance, ROC and PR curves for RBFOX2

Additional File 2: Classifier Metrics and Results for all 112 RBPs

Additional File 3: K-mer enrichment scores and percentages Additional File 4: Overlap percentages

Additional File 5: Structure percentages

Additional File 6: Heatmap with Z-score for binding region preference

Additional File 7: RBP classification for Kmer/Struct/Overlap and combinations

Additional File 8: List of accession numbers for RBPs from ENCODE

Additional File 9: BioFeatureFinder v1.0 algorithm

## References

1. Reuter JA, Spacek D V., Snyder MP. High-Throughput Sequencing Technologies. Mol. Cell. 2015. p. 586–97.

2. Park PJ. ChIP-seq: Advantages and challenges of a maturing technology. Nat. Rev. Genet. 2009. p. 669–80.

3. Ule J, Jensen KB, Ruggiu M, Mele A, Ule A, Darnell RB. CLIP Identifies Nova-Regulated RNA Networks in the Brain. Science (80-.). 2003;302:1212–5.

4. Pan Q, Shai O, Lee LJ, Frey BJ, Blencowe BJ. Deep surveying of alternative splicing complexity in the human transcriptome by high-throughput sequencing. Nat. Genet. 2008;40:1413–5.

5. Rapaport F, Khanin R, Liang Y, Pirun M, Krek A, Zumbo P, et al. Comprehensive evaluation of differential gene expression analysis methods for RNA-seq data. Genome Biol. 2013;14.

6. Kircher M, Kelso J. High-throughput DNA sequencing--concepts and limitations. Bioessays [Internet]. 2010;32:524–36. Available from: http://dx.doi.org/10.1002/bies.200900181%5 http://onlinelibrary.wiley.com/store/10.1002/bies.200900181/asset/524_ftp.pdf?v=1&t=hz4sgjb3&s=d7669c48bf7843ed9e0715d320188c85879897ca

7. Huber W, Carey VJ, Gentleman R, Anders S, Carlson M, Carvalho BS, et al. Orchestrating high-throughput genomic analysis with Bioconductor. Nat. Methods. 2015;12:115–21.

8. Conesa A, Madrigal P, Tarazona S, Gomez-Cabrero D, Cervera A, McPherson A, et al. A survey of best practices for RNA-seq data analysis. Genome Biol. [Internet]. 2016;17:13. Available from: http://www.ncbi.nlm.nih.gov/pubmed/26813401%5 http://www.pubmedcentral.nih.gov/articlerender.fcgi?artid=PMC4728800

9. Uhl M, Houwaart T, Corrado G, Wright PR, Backofen R. Computational analysis of CLIP-seq data. Methods. 2017. p. 60–72.

10. Liu Q, Zhong X, Madison BB, Rustgi AK, Shyr Y. Assessing Computational Steps for CLIP-Seq Data Analysis. Biomed Res. Int. 2015;2015.

11. Steinhauser S, Kurzawa N, Eils R, Herrmann C. A comprehensive comparison of tools for differential ChIP-seq analysis. Brief. Bioinform. [Internet]. 2016;bbv110. Available from: https://academic.oup.com/bib/article-lookup/doi/10.1093/bib/bbv110

12. Bailey T, Krajewski P, Ladunga I, Lefebvre C, Li Q, Liu T, et al. Practical Guidelines for the Comprehensive Analysis of ChIP-seq Data. PLos Comput. Biol. 2013;9.

13. Maticzka D, Lange SJ, Costa F, Backofen R. GraphProt: modeling binding preferences of RNA-binding proteins. Genome Biol. [Internet]. 2014;15:R17. Available from: http://www.pubmedcentral.nih.gov/articlerender.fcgi?artid=4053806&tool=pmcentrez&rendertype=abstract

14. Zhang MQ. Statistical features of human exons and their flanking regions. Hum. Mol. Genet. 1998;7:919–32.

15. Pesole G, Mignone F, Gissi C, Grillo G, Licciulli F, Liuni S. Structural and functional features of eukaryotic mRNA untranslated regions. Gene. 2001. p. 73–81.

16. Zhang MQ. Computational prediction of eukaryotic protein-coding genes. Nat. Rev. Genet. 2002. p. 698–709.

17. Larsen F, Gundersen G, Lopez R, Prydz H. CpG islands as gene markers in the human genome. Genomics. 1992;13:1095–107.

18. Siepel A, Bejerano G, Pedersen JS, Hinrichs AS, Hou M, Rosenbloom K, et al. Evolutionarily conserved elements in vertebrate, insect, worm, and yeast genomes. Genome Res. 2005;15:1034–50.

19. Sethupathy P, Collins FS. MicroRNA target site polymorphisms and human disease. Trends Genet. 2008;24:489–97.

20. Reczko M, Maragkakis M, Alexiou P, Grosse I, Hatzigeorgiou AG. Functional microRNA targets in protein coding sequences. Bioinformatics [Internet]. 2012;28:771–6. Available from: http://www.ncbi.nlm.nih.gov/pubmed/22285563

21. Yip KY, Cheng C, Bhardwaj N, Brown JB, Leng J, Kundaje A, et al. Classification of human genomic regions based on experimentally determined binding sites of more than 100 transcription-related factors. Genome Biol. 2012;13.

22. Schübeler D. Function and information content of DNA methylation. Nature. 2015. p. 321–6.

23. Raychaudhuri S, Plenge RM, Rossin EJ, Ng ACY, Purcell SM, Sklar P, et al. Identifying relationships among genomic disease regions: Predicting genes at pathogenic SNP associations and rare deletions. Plos Genet. 2009;5.

24. Subramanian S, Mishra RK, Singh L. Genome-wide analysis of microsatellite repeats in humans: their abundance and density in specific genomic regions. Genome Biol. 2003;4:R13.

25. 1000 Genomes Project Consortium T 1000 GP, Abecasis GR, Auton A, Brooks LD, DePristo MA, Durbin RM, et al. An integrated map of genetic variation from 1,092 human genomes. Nature [Internet]. 2012;491:56–65. Available from: http://www.ncbi.nlm.nih.gov/pubmed/23128226%5 http://www.pubmedcentral.nih.gov/articlerender.fcgi?artid=PMC3498066

26. Korbel JO, Urban AE, Affourtit JP, Godwin B, Grubert F, Simons JF, et al. Paired-End Mapping Reveals Extensive Structural Variation in the Human Genome. Science (80-.). [Internet]. 2007;318:420–6. Available from: http://www.sciencemag.org/cgi/doi/10.1126/science.1149504

27. Stein L. Genome annotation: From sequence to biology. Nat. Rev. Genet. 2001. p. 493–503.

28. Liu C, Che D, Liu X, Song Y. Applications of machine learning in genomics and systems biology. Comput. Math. Methods Med. 2013;2013.

29. Libbrecht MW, Noble WS. Machine learning in genetics and genomics. 2017;16:321–32.

30. Zhang YQ, Rajapakse JC. Machine Learning in Bioinformatics. Mach. Learn. Bioinforma. 2008.

31. Singireddy S, Alkhateeb A, Rezaeian I, Rueda L, Cavallo-Medved D, Porter L. Identifying differentially expressed transcripts associated with prostate cancer progression using RNA-Seq and machine learning techniques. 2015 IEEE Conf. Comput. Intell. Bioinforma. Comput. Biol. [Internet]. 2015. p. 1–5. Available from: http://ieeexplore.ieee.org/document/7300302/

32. Xue B, Oldfield CJ, Dunker AK, Uversky VN. CDF it all: Consensus prediction of intrinsically disordered proteins based on various cumulative distribution functions. FEBS Lett. 2009;583:1469–74.

33. Kent WJ, Sugnet CW, Furey TS, Roskin KM, Pringle TH, Zahler AM, et al. The Human Genome Browser at UCSC. Genome Res. 2002. p. 996–1006.

34. Yates A, Akanni W, Amode MR, Barrell D, Billis K, Carvalho-Silva D, et al. Ensembl 2016. Nucleic Acids Res. 2016;44:D710–6.

35. Harrow J, Frankish A, Gonzalez JM, Tapanari E, Diekhans M, Kokocinski F, et al. GENCODE: The reference human genome annotation for the ENCODE project. Genome Res. 2012;22:1760–74.

36. Encode Consortium. An integrated encyclopedia of DNA elements in the human genome. Nature. 2013;489:57–74.

37. Lovci MT, Ghanem D, Marr H, Arnold J, Gee S, Parra M, et al. Rbfox proteins regulate alternative mRNA splicing through evolutionarily conserved RNA bridges. Nat. Struct. Mol. Biol. 2013;20:1434–42.

38. Wei C, Xiao R, Chen L, Cui H, Zhou Y, Xue Y, et al. RBFox2 Binds Nascent RNA to Globally Regulate Polycomb Complex 2 Targeting in Mammalian Genomes. Mol. Cell. 2016;62:875–89.

39. Van Nostrand EL, Pratt GA, Shishkin AA, Gelboin-Burkhart C, Fang MY, Sundararaman B, et al. Robust transcriptome-wide discovery of RNA-binding protein binding sites with enhanced CLIP (eCLIP). Nat. Methods [Internet]. 2016;13:1–9. Available from: http://www.nature.com/doifinder/10.1038/nmeth.3810%5 http://www.ncbi.nlm.nih.gov/pubmed/27018577

40. Jangi M, Boutz PL, Paul P, Sharp PA. Rbfox2 controls autoregulation in RNA-binding protein networks. Genes Dev. 2014;28:637–51.

41. Quinlan AR, Hall IM. BEDTools: A flexible suite of utilities for comparing genomic features. Bioinformatics. 2010;26:841–2.

42. Kent WJ, Zweig AS, Barber G, Hinrichs AS, Karolchik D. BigWig and BigBed: Enabling browsing of large distributed datasets. Bioinformatics. 2010;26:2204–7.

43. Rice P, Longden I, Bleasby A. EMBOSS: The European Molecular Biology Open Software Suite. Trends Genet. [Internet]. 2000;16:276–7. Available from: http://linkinghub.elsevier.com/retrieve/pii/S0168952500020242

44. Tafer H, Höner zu Siederdissen C, Stadler PF, Bernhart SH, Hofacker IL, Lorenz R, et al. ViennaRNA Package 2.0. Algorithms Mol. Biol. 2011;6:26.

45. Kikin O, D’Antonio L, Bagga PS. QGRS Mapper: A web-based server for predicting G-quadruplexes in nucleotide sequences. Nucleic Acids Res. 2006;34.

46. Oliphant TE. SciPy: Open source scientific tools for Python. Comput. Sci. Eng. [Internet]. 2007;9:10–20. Available from: http://www.scipy.org/

47. Weston J, Mukherjee S, Chapelle O, Pontil M. Feature selection for SVMs. Nips [Internet]. 2000;13:668–74. Available from: http://www.ee.columbia.edu/∼sfchang/course/svia-F04/slides/feature_selection_for_SVMs.pdf

48. Ivanov A, Riccardi G. Kolmogorov-Smirnov test for feature selection in emotion recognition from speech. ICASSP, IEEE Int. Conf. Acoust. Speech Signal Process. -Proc. 2012. p. 5125–8.

49. Subrahmanyam K, Sankar NS, Baggam SP. A Modified KS-test for Feature Selection. IOSR J. Comput. Eng. 2013;13:73–9.

50. Pedregosa F, Varoquaux G, Gramfort A, Michel V, Thirion B, Grisel O, et al. Scikit-learn: Machine Learning in Python. J. Mach. Learn. Res. [Internet]. 2012;12:2825–30. Available from: http://dl.acm.org/citation.cfm?id=207819/ http://arxiv.org/abs/1201.0490

51. Blagus R, Lusa L. Boosting for High-Dimensional Two-Class Prediction. BMC Bioinformatics [Internet]. 2015;16:300. Available from: http://www.biomedcentral.com/1471-2105/16/300

52. Galar M, Fernandez A, Barrenechea E, Bustince H, Herrera F. A review on ensembles for the class imbalance problem: Bagging-, boosting-, and hybrid-based approaches. IEEE Trans. Syst. Man Cybern. Part C Appl. Rev. 2012. p. 463–84.

53. Schapire RE. A brief introduction to boosting. IJCAI Int. Jt. Conf. Artif. Intell. 1999. p. 1401–6.

54. Hastie T, Tibshirani R, Friedman J. Relative Importance of Predictor Variables. Elem. Stat. Learn. Elem. Stat. Learn. Mining, Inference, Predict. Second Ed. [Internet]. 2001. p. 367–9. Available from: http://www.springerlink.com/index/D7X7KX6772HQ2135.pdf%255 http://www-stat.stanford.edu/∼tibs/book/preface.ps

55. Longadge R, Dongre SS, Malik L. Class imbalance problem in data mining: review. Int. J. Comput. Sci. Netw. 2013;2:83–7.

56. Chan CY, Carmack CS, Long DD, Maliyekkel A, Shao Y, Roninson IB, et al. A structural interpretation of the effect of GC-content on efficiency of RNA interference. BMC Bioinformatics. 2009;10:1–7.

57. Zhang J, Kuo CCJ, Chen L. GC content around splice sites affects splicing through pre-mRNA secondary structures. BMC Genomics [Internet]. 2011;12:90. Available from: http://www.biomedcentral.com/1471-2164/12/90

58. Li X, Kazan H, Lipshitz HD, Morris QD. Finding the target sites of RNA-binding proteins. Wiley Interdiscip. Rev. RNA. 2014. p. 111–30.

59. Kazan H, Ray D, Chan ET, Hughes TR, Morris Q. RNAcontext: A new method for learning the sequence and structure binding preferences of RNA-binding proteins. Plos Comput. Biol. 2010;6:28.

60. Thandapani P, O’Connor TR, Bailey TL, Richard S. Defining the RGG/RG Motif. Mol. Cell. 2013;50:613–23.

61. Vasilyev N, Polonskaia A, Darnell JC, Darnell RB, Patel DJ, Serganov A. Crystal structure reveals specific recognition of a G-quadruplex RNA by a β-turn in the RGG motif of FMRP. Proc. Natl. Acad. Sci. U. S. A. [Internet]. 2015;112:E5391–400. Available from: http://www.ncbi.nlm.nih.gov/pubmed/26374839%5 http://www.pubmedcentral.nih.gov/articlerender.fcgi?artid=PMC4593078

62. Marcel V, Tran PLT, Sagne C, Martel-Planche G, Vaslin L, Teulade-Fichou MP, et al. G-quadruplex structures in TP53 intron 3: Role in alternative splicing and in production of p53 mRNA isoforms. Carcinogenesis. 2011;32:271–8.

63. Ribeiro MM, Teixeira GS, Martins L, Marques MR, AP de Souza, Line SRP. G-quadruplex formation enhances splicing efficiency of PAX9 intron 1. Hum. Genet. 2014;134:37–44.

64. Cary GA, Vinh DBN, May P, Kuestner R, Dudley AM. Proteomic Analysis of Dhh1 Complexes Reveals a Role for Hsp40 Chaperone Ydj1 in Yeast P-Body Assembly. G3; Genes|Genomes|Genetics [Internet]. 2015;5:2497–511. Available from: http://g3journal.org/lookup/doi/10.1534/g3.115.021444

65. Chatr-Aryamontri A, Oughtred R, Boucher L, Rust J, Chang C, Kolas NK, et al. The BioGRID interaction database: 2017 update. Nucleic Acids Res. 2017;45:D369–79.

66. Szklarczyk D, Morris JH, Cook H, Kuhn M, Wyder S, Simonovic M, et al. The STRING database in 2017: Quality-controlled protein-protein association networks, made broadly accessible. Nucleic Acids Res. 2017;45:D362–8.

67. Huttlin EL, Ting L, Bruckner RJ, Gebreab F, Gygi MP, Szpyt J, et al. The BioPlex Network: A Systematic Exploration of the Human Interactome. Cell. 2015;162:425–40.

68. Huttlin EL, Bruckner RJ, Paulo JA, Cannon JR, Ting L, Baltier K, et al. Architecture of the human interactome defines protein communities and disease networks. Nature. 2017;545:505–9.

69. Lim J, Hao T, Shaw C, Patel AJ, Szabó G, Rual JF, et al. A Protein-Protein Interaction Network for Human Inherited Ataxias and Disorders of Purkinje Cell Degeneration. Cell. 2006;125:801–14.

70. Hegele A, Kamburov A, Grossmann A, Sourlis C, Wowro S, Weimann M, et al. Dynamic Protein-Protein Interaction Wiring of the Human Spliceosome. Mol. Cell. 2012;45:567–80.

71. Havugimana PC, Hart GT, Nepusz T, Yang H, Turinsky AL, Li Z, et al. A census of human soluble protein complexes. Cell. 2012;150:1068–81

72. Shao C, Yang B, Wu T, Huang J, Tang P, Zhou Y, et al. Mechanisms for U2AF to define 3â €2 splice sites and regulate alternative splicing in the human genome. Nat. Struct. Mol. Biol. 2014;21:997–1005.

73. Tanackovic G, Krämer A. Human splicing factor SF3a, but not SF1, is essential for pre-mRNA splicing in vivo. Mol. Biol. Cell [Internet]. 2005;16:1366– Available from: http://www.pubmedcentral.nih.gov/articlerender.fcgi?artid=551499&tool=pmcentrez&rendertype=abstract

74. Wickramasinghe VO, Gonzàlez-Porta M, Perera D, Bartolozzi AR, Sibley CR, Hallegger M, et al. Regulation of constitutive and alternative mRNA splicing across the human transcriptome by PRPF8 is determined by 5’ splice site strength. Genome Biol. 2015;16.

75. Ascano M, Mukherjee N, Bandaru P, Miller JB, Nusbaum JD, Corcoran DL, et al. FMRP targets distinct mRNA sequence elements to regulate protein expression. Nature. 2012;492:382–6.

76. Lu G, Hall TMT. Alternate modes of cognate RNA recognition by human PUMILIO proteins. Structure. 2011;19:361–7.

77. Matoulkova E, Michalova E, Vojtesek B, Hrstka R. The role of the 3′ untranslated region in post-transcriptional regulation of protein expression in mammalian cells. RNA Biol. 2012. p. 563–76.

78. Tollervey JR, Curk T, Rogelj B, Briese M, Cereda M, Kayikci M, et al. Characterizing the RNA targets and position-dependent splicing regulation by TDP-43. Nat. Neurosci. [Internet]. Nature Publishing Group; 2011 [cited 2014 Mar 22];14:452–8. Available from: http://www.pubmedcentral.nih.gov/articlerender.fcgi?artid=3108889&tool=pmcentrez&rendertype=abstract

79. Lou T-F, Weidmann CA, Killingsworth J, Tanaka Hall TM, Goldstrohm AC, Campbell ZT. Integrated analysis of RNA-binding protein complexes using in vitro selection and high-throughput sequencing and sequence specificity landscapes (SEQRS). Methods [Internet]. Elsevier Inc.; 2017;118-119:171–81. Available from: http://linkinghub.elsevier.com/retrieve/pii/S1046202316303383

80. Campbell ZT, Wickens M. Probing RNA-protein networks: Biochemistry meets genomics. Trends Biochem. Sci. [Internet]. Elsevier Ltd; 2015;40:157–64. Available from: http://dx.doi.org/10.1016/j.tibs.2015.01.003

81. Siomi MC, Zhang Y, Siomi H, Dreyfuss G. Specific sequences in the fragile X syndrome protein FMR1 and the FXR proteins mediate their binding to 60S ribosomal subunits and the interactions among them. Mol. Cell. Biol. [Internet]. 1996;16:3825–32. Available from: http://mcb.asm.org/lookup/doi/10.1128/MCB.16.7.3825

82. Colombrita C, Onesto E, Megiorni F, Pizzuti A, Baralle FE, Buratti E, et al. TDP-43 and FUS RNA-binding proteins bind distinct sets of cytoplasmic messenger RNAs and differently regulate their post-transcriptional fate in motoneuron-like cells. J. Biol. Chem. 2012;287:15635–47.

83. Galarneau A, Richard S. Target RNA motif and target mRNAs of the Quaking STAR protein. Nat. Struct. Mol. Biol. 2005;12:691–8.

84. Pérez I, Lin CH, McAfee JG, Patton JG. Mutation of PTB binding sites causes misregulation of alternative 3’ splice site selection in vivo. RNA [Internet]. 1997;3:764–78. Available from: http://rnajournal.cshlp.org/content/3/7/764.abstract

85. König J, Zarnack K, Rot G, Curk T, Kayikci M, Zupan B, et al. ICLIP reveals the function of hnRNP particles in splicing at individual nucleotide resolution. Nat. Struct. Mol. Biol. 2010;17:909–15.

86. Swanson MS, Dreyfuss G. Classification and purification of proteins of heterogeneous nuclear ribonucleoprotein particles by RNA-binding specificities. Mol. Cell. Biol. [Internet]. 1988;8:2237–41. Available from: http://www.pubmedcentral.nih.gov/articlerender.fcgi?artid=363409&tool=pmcentrez&rendertype=abstract

87. Galarneau A, Richard S. The STAR RNA binding proteins GLD-1, QKI, SAM68 and SLM-2 bind bipartite RNA motifs. BMC Mol. Biol. 2009;10.

88. Dember LM, Kim ND, Liu KQ, Anderson P. Individual RNA recognition motifs of TIA-1 and TIAR have different RNA binding specificities. J. Biol. Chem. 1996;271:2783–8.

89. Takahama K, Kino K, Arai S, Kurokawa R, Oyoshi T. Identification of Ewing’s sarcoma protein as a G-quadruplex DNA-and RNA-binding protein. FEBS J. 2011;278:988–98.

90. Skourti-Stathaki K, Proudfoot NJ, Gromak N. Human Senataxin Resolves RNA/DNA Hybrids Formed at Transcriptional Pause Sites to Promote Xrn2-Dependent Termination. Mol. Cell. 2011;42:794–805.

91. Stefanovic S, DeMarco BA, Underwood A, Williams KR, Bassell GJ, Mihailescu MR. Fragile X mental retardation protein interactions with a G quadruplex structure in the 3’-untranslated region of NR2B mRNA. Mol. Biosyst. [Internet]. 2015;11:3222–30. Available from: http://www.pubmedcentral.nih.gov/articlerender.fcgi?artid=4643373&tool=pmcentrez&rendertype=abstract

92. Cammas A, Millevoi S. RNA G-quadruplexes: emerging mechanisms in disease. Nucleic Acids Res. 2017;45:1584–95.

93. BioFeatureFindder [Internet]. GitHub. Available from: https://github.com/kbmlab/BioFeatureFinder

