## Supplementary figures and images for "BioFeatureFinder: Flexible, unbiased analysis of biological characteristics associated with genomic regions"

### Supplementary Materials

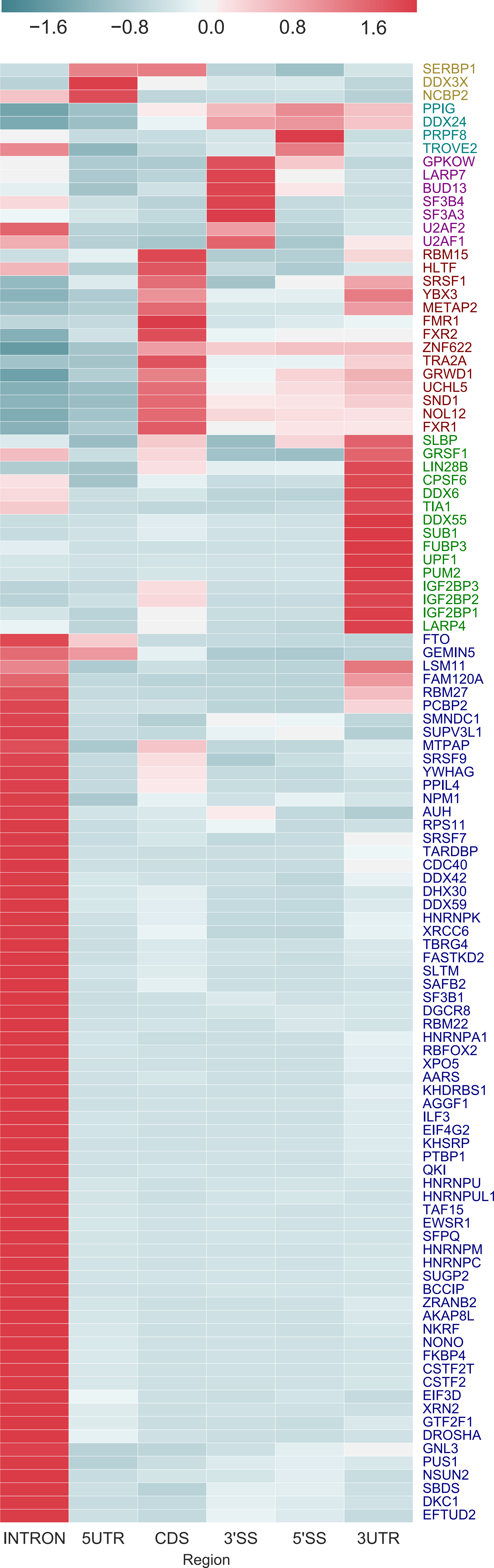

### Supplementary Materials

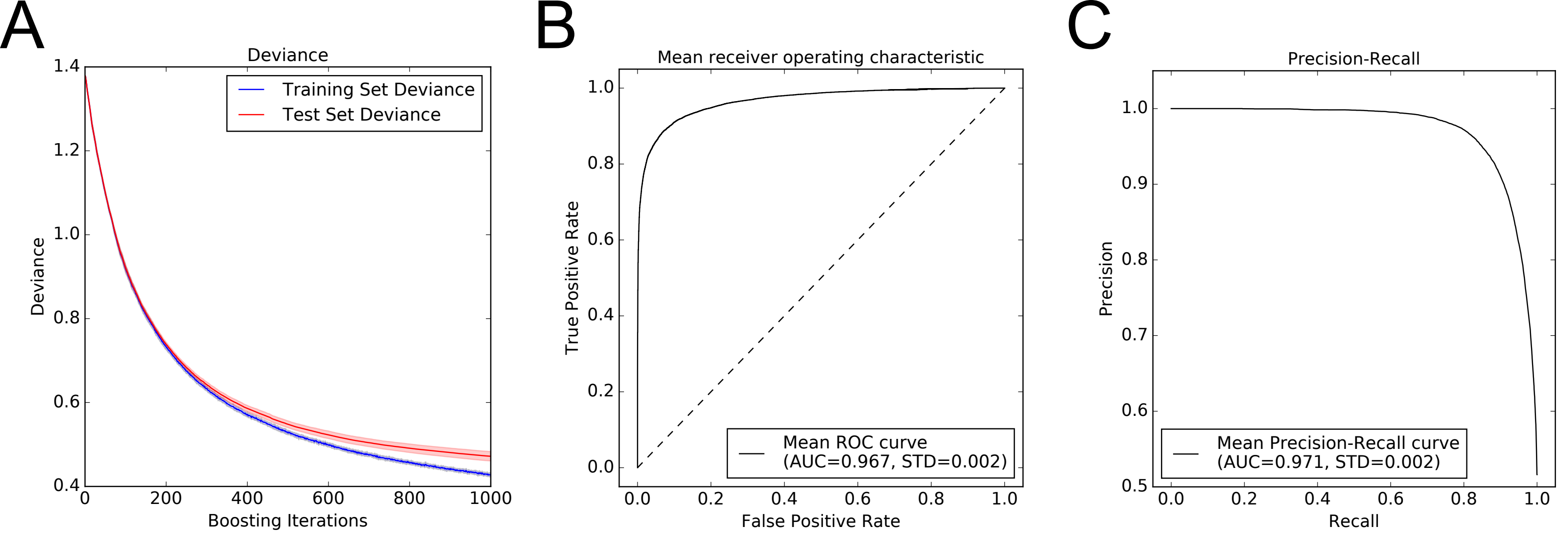

### Supplementary Materials

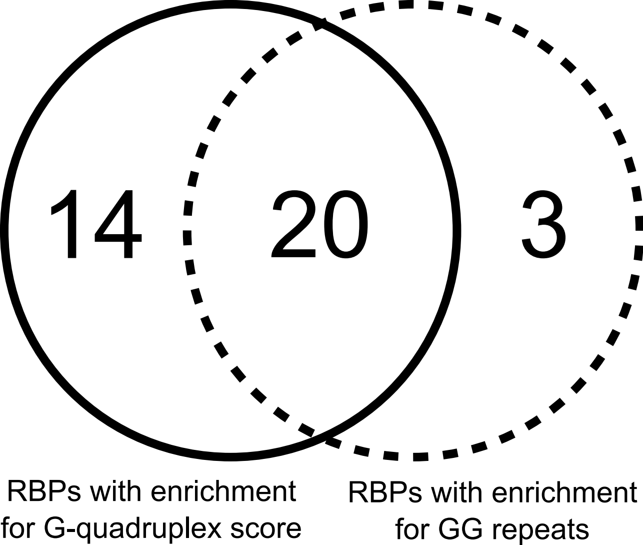
